# Collaboration Between Host and Viral Factors Shape SARS-CoV-2 Evolution

**DOI:** 10.1101/2021.07.16.452629

**Authors:** Connor G. G. Bamford, Lindsay Broadbent, Elihu Aranday-Cortes, Mary McCabe, James McKenna, David Courtney, Olivier Touzelet, Ahlam Ali, Grace Roberts, Guillermo Lopez Campos, David Simpson, Conall McCaughey, Derek Fairley, Ken Mills, Ultan F. Power, the Breathing Together Investigators

## Abstract

SARS-CoV-2 continues to evolve, resulting in several ‘variants of concern’ with novel properties. The factors driving SARS-CoV-2 fitness and evolution in the human respiratory tract remain poorly defined. Here, we provide evidence that both viral and host factors co-operate to shape SARS-CoV-2 genotypic and phenotypic change. Through viral whole-genome sequencing, we explored the evolution of two clinical isolates of SARS-CoV-2 during passage in unmodified Vero-derived cell lines and in parallel, in well-differentiated primary nasal epithelial cell (WD-PNEC) cultures. We identify a consistent, rich genetic diversity arising in vitro, variants of which could rapidly rise to near-fixation with 2 passages. Within isolates, SARS-CoV-2 evolution was dependent on host cells, with Vero-derived cells facilitating more profound genetic changes. However, most mutations were not shared between strains. Furthermore, comparison of both Vero-grown isolates on WD-PNECs disclosed marked growth attenuation mapping to the loss of the polybasic cleavage site (PBCS) in Spike while the strain with mutations in NSP12 (T293I), Spike (P812R) and a truncation of ORF7a remained viable in WD-PNECs. Our work highlights the significant genetic plasticity of SARS-CoV-2 while uncovering an influential role for collaboration between viral and host cell factors in shaping viral evolution and fitness in human respiratory epithelium.

## Introduction

Severe acute respiratory syndrome coronavirus 2 (SARS-CoV-2) (Family: *Coronaviridae*; Genus: *Betacoronavirus*) has a positive-sense, non-segmented, single-stranded RNA genome of ~30,000 nucleotides in length (**Lu et al., 2020; Wu et al., 2020**). The SARS-CoV-2 genome encodes at least 29 proteins, expressed from translation of a 5’ major open reading frame (ORF1ab), including Nsp3 and Nsp12 (viral RNA-dependent RNA polymerase), and a series of nested transcripts at the 3’ terminus, including Spike (S; the viral attachment and fusion glycoprotein) and ORF7a. SARS-CoV-2 emerged into the human population in late 2019, causing coronavirus virus disease 2019 (COVID-19) (**Wang et al., 2020**). Reflecting its likely zoonotic origins, SARS-CoV-2-like and other SARS-related viruses have been detected and isolated from horseshoe bats and pangolins from Asia (Boni et al., 2020). SARS-CoV-2 is a highly transmissible virus with an R0 of up to ~5, and has a relatively high case mortality rate (~1%), especially pathogenic in elderly or individuals with co-morbidities (**Cevik et al., 2020**). While safe and effective vaccines were recently developed (**Krammer, 2020**), there is a dearth of highly-effective therapeutic interventions, with notable exceptions such as dexamethasone (**Recovery Group, 2021**).

SARS-CoV-2 productively infects the epithelial cells lining the upper and lower respiratory tract, including those in the nasal cavity and the alveoli of the lung (**Hou et al., 2020**). By virtue of interaction with Spike, SARS-CoV-2 exploits host cell protein angiotensin-converting enzyme 2 (ACE2) as its receptor (**Shang et al., 2020**). Additionally, for entry to occur Spike requires activation by two host proteases, furin and transmembrane protease serine 2 (TMPRSS2)-like proteases, which cleave Spike at the S1/S2 boundary between its two subunits (S1 and S2) and the S2’ site in S2 allowing release of the fusion peptide (**Hoffmann et al., 2020**). Following binding to ACE2, a proteolytically-activated Spike can fuse the viral envelope with the host cell membrane releasing the infectious genome into the cytoplasm.

Since its initial emergence, SARS-CoV-2 has continued to evolve and adapt to the human population with several putatively beneficial mutations arising in Spike, such as D614G and N501Y, that affect Spike stability and binding to ACE2, and antibody-escape mutations in the amino-terminal domain (NTD) (**Harvey et al., 2021**). Additionally, loss of the polybasic cleavage site (PBCS), which is a unique feature of SARS-CoV-2 and facilitates furin cleavage at the S1/S2 boundary, has been demonstrated to reduce transmission and virulence of SARS-CoV-2 in animal models (**Johnson et al., 2021; Peacock et al., 2021**). Together, these mutations of interest are found in constellations in so-called ‘variants of concern’ (VOC), which are strains of SARS-CoV-2 with evident phenotypic differences, such as enhanced transmissibility, pathogenicity, or reduced sensitivity to antibody-mediated neutralization in humans (**Harvey et al., 2021**).

As SARS-CoV-2 continues to spread, and interventions and vaccines are being rolled out, there remain significant unknowns as to how SARS-CoV-2 may adapt further to humans. Knowledge of the genetic and molecular correlates of this difference in transmissibility is crucial for understanding of coronavirus pandemic preparedness and inform strategies for surveillance and control. In vitro models can help disentangle the factors affecting evolution, identify new ones, and highlight mutational tolerance. Here, we undertook a side-by-side comparison of SARS-CoV-2 evolution by whole genome sequencing of two isolates, grown in parallel in standard Vero-derived cells and human ‘well-differentiated primary nasal epithelial cells’ (WD-PNECs), which are a useful model for probing virus-host interactions in the respiratory tract (**Guo-Parke et al., 2013; Hou et al., 2020; Villenave et al., 2012**). Our data demonstrate clear roles of both viral and host cell factors in shaping SARS-CoV-2 genetic and functional changes, identifying genetic features required for efficient infection of primary cells.

## Results

### Isolation and passage of SARS-CoV-2 in unmodified Vero-derived cells

To begin to understand the evolution of SARS-CoV-2 we first needed to generate characterised stocks of virus (**Fig 1A**). In the first instance, a low passage isolate (passage 1, P1) of SARS-CoV-2 (England 02/20) was obtained from Public Health England (PHE) and is referred to as ‘PHE’. This stock was from a sample isolated on VeroE6 cells and represents one of the earliest isolates of SARS-CoV-2 in the UK during the pandemic. PHE is from clade A and does not contain the D614G substitution in Spike (**supplementary table 1**)(**Davidson et al., 2020; Holden et al., 2020**). Upon receipt, we carried out a further three passages on VeroE6 cells passaging at an MOI of 0.001, harvesting stocks at 96 hpi when extensive cytopathic effect was observed. The PHE strain grew efficiently, reaching titres of >10^6^ pfu/mL (**Fig 1B**) and was cytopathic, inducing ‘webbing’ and cell rounding, consistent with previous reports (data not shown).

**Figure 1.**
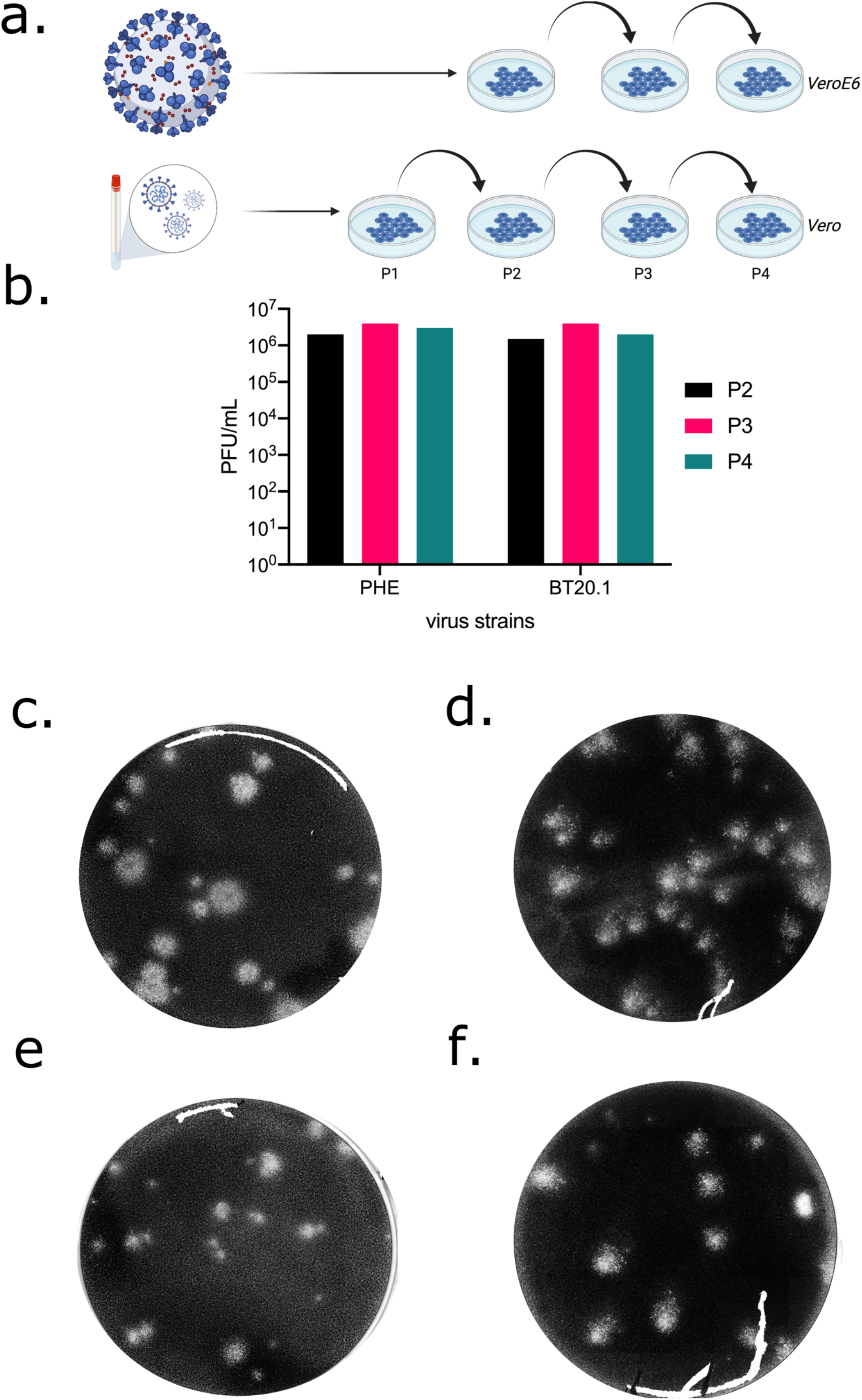
Isolation and passage of PHE and BT20.1 in Vero-derived cells. Schematic of SARS-CoV-2 isolation/serial passage series on VeroE6 or Vero cells for PHE and BT20.1 from isolation to P4 (a). Extracellular infectivity titres for stocks generated from P2-P4 VeroE6/Vero passage for PHE and BT20.1 using plaque assay protocol on Vero cells (b). Plaque visualisation of PHE (c) and BT20.1 (d) P4, and P2 (e and f) on Vero cells. Figures were generated with the aid of Biorender (https://biorender.com/).

As we wanted to understand the viral factors that may drive evolution and results obtained from only one isolate may be non-representative, we next obtained an independent – but comparable - clinical nasal/pharyngeal swab sample containing SARS-CoV-2, which we termed BT20.1 (Belfast 06/20). This strain represents an isolate from the UK’s ‘first wave’ and is a representative of clade B that contains the D614G mutation in Spike (table 1). Unlike PHE, BT20.1 was isolated on standard Vero cells (CCL-81) and passaged to P4 (multiplicity of infection [MOI]~0.001 and passaged every 3 days). Like PHE, BT20.1 grew efficiently, reaching titres of >10^6^ pfu/mL (**Fig 1B**).

Both PHE and BT20.1 formed plaques on standard Vero cells in all passages (**Fig 1C and D**). Comparison of plaque sizes between P2 and P4 identified differences in plaque size composition following Vero cell passage. This was most evident for PHE, which became predominantly large plaques (**Fig 1 E and F**). As observed in the plaque edge, BT20.1 induced consistent cell-to-cell fusion, unlike PHE (**sFig 1 A and B**).

### Sequencing of SARS-CoV-2 passage series in unmodified Vero-derived cells

As we had successfully generated comparable in vitro passage series for two relevant isolates of SARS-CoV-2, we next determined what genetic changes, if any, occurred during passage in unmodified Vero-derived cells. Whole genome sequences of our SARS-CoV-2 stocks at each passage were generated and minor sequence analysis (>5% minor allele frequency) was carried out, comparing variation arising to the Wuhan-Hu-1 reference (NC_045512.2) genome sequence for SARS-CoV-2 (**supplementary table 1**). Unfortunately, the sequence depth and quality were not sufficient to reconstruct a whole genome sequences for BT20.1 P1 isolate material, likely due to insufficient viral material resulting from the initial isolation. Therefore, we focused our analysis on PHE P1-4 and BT20.1 P2-P4.

Analysing mutations in the PHE passage series we identified 4 changes (C8782T; T18488T; T28144C; A29596G) relative to Wuhan-Hu-1 consistently at ~100% at all passages, likely reflecting fixation in the original virus stock (**Fig 2A**). These changes were considered intrinsic to that particular strain and were not analysed any further herein as we wished to focus on variants arising during passage. Sequencing confirmed the presence of D614 in Spike, consistent with it being an early SARS-CoV-2 isolate.

**Figure 2.**
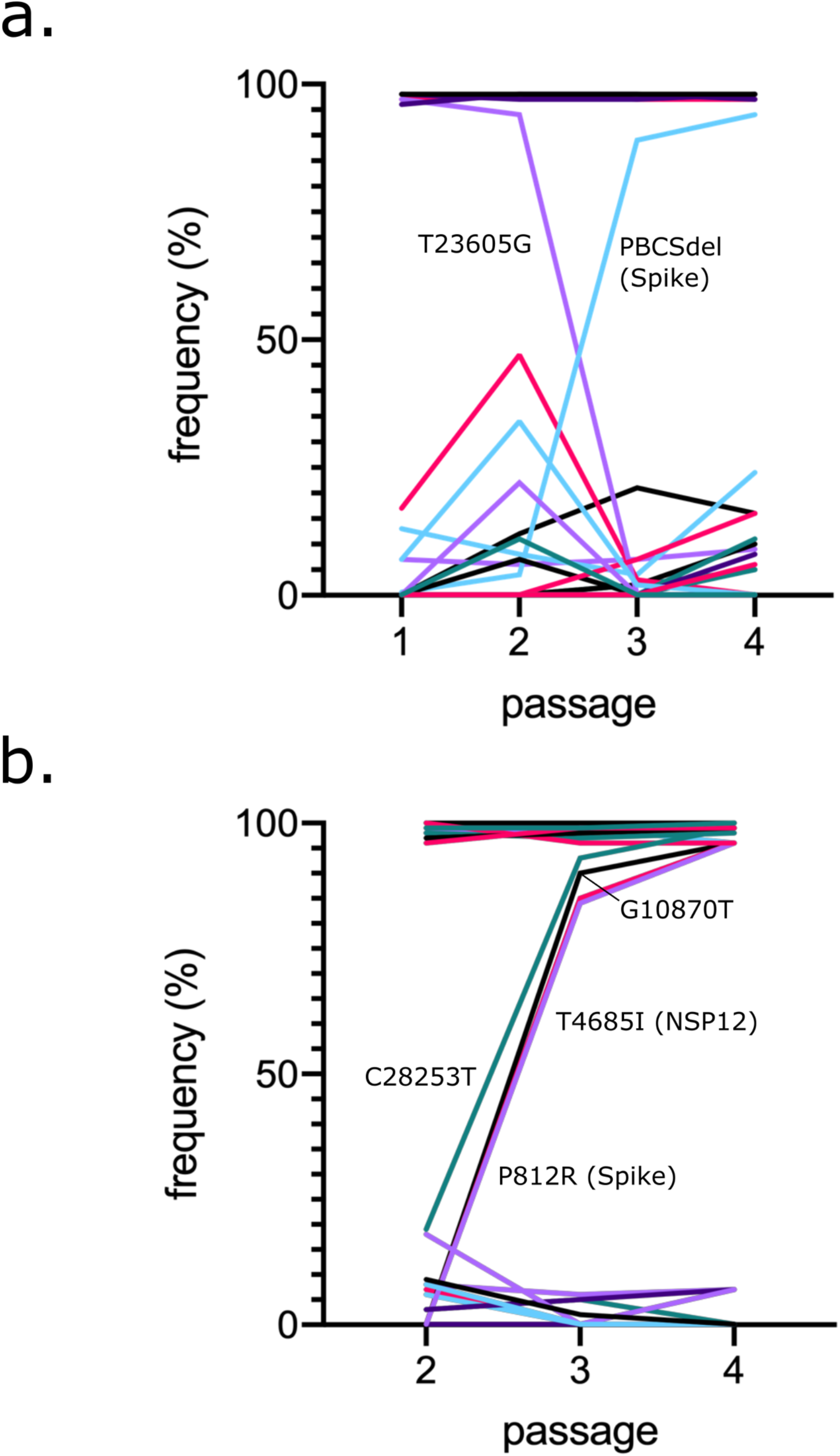
Analysis of PHE and BT20.1 whole genome sequences during Vero cell passage. Frequency of mutations detected for PHE (a) and BT20.1 (b) passage series on VeroE6 or Vero cells, respectively, relative to the reference sequence (Wuhan-Hu-1). Only sequences from P1-P4 (PHE) and P2-P4 (BT20.1) were analysed to facilitate adequate comparisons. Core changes are found consistently at high frequency and minor variants found at consistently low frequency (e.g. <50%). Only variants that significantly changed in frequency are marked on the graph. Colours do not reflect relationships between variants.

Outwith the core changes described above, two major mutations were observed: a synonymous (T23605G) and non-synonymous out-of-frame deletion (deletion of 24 nucleotides AATTCTCCTCGGCGGGCACGTAGTG 23597A; resulting in the replacement of of 9 amino acids [679-687; NSPRRARSV] in Spike with an isoleucine [I]) mapping to the polybasic cleavage site (PBCS) (**Fig 2A**). Deletion of the PBCS ablated the T23605G synonymous variant in the process. This occurred at P3, although the deletion was observed in the original P1 material from PHE. Furthermore, we detected 15 minor variants (non-consensus) that had an allele frequency (AF) of >5% in at least one sample of the passage series. These changes mapped to several genes and proteins of SARS-CoV-2, including ORF1AB, Spike, E, N, and ORF10. (**supplementary table 1**). Interestingly, we observed a cluster of three mutations occurring in the amino terminal domain (NTD) of Spike, appearing at P3 and rising in frequency at P4. Two of these Spike NTD mutations were similar to mutations occurring in VOCs: D215G and a deletion of 24 nucleotides (GCTATACATGTCTCTGGGACCAATGGTA21761G) resulting in a loss of 9 amino acids IHVSGTNGT (aa68-77). Additionally, we noticed a convergent mutation of L37 in E, detecting two mutations resulting in L37F and L37R. To determine the reproducibility of passage sequencing, an independent P4 PHE (P4B) was generated from P3 and sequenced, with very high levels of similarity between the two (**supplementary table 1**).

Like PHE, we identified core changes inherent to BT20.1 (**Fig 2B**), which were greater in number than PHE (10 vs 4), consistent with its later isolation (February 2020 versus June 2020) (**supplementary table 1**). These changes included, but were not limited to, D614G in Spike; R203K & G204K in N; and an out-of-frame deletion of 5 nucleotides in ORF7A leading to its premature truncation. Like PHE, we identified mutations arising rapidly upon consecutive passage in Vero cells (i.e., were not detected at P2), including the non-synonymous mutations T293I in NSP12 and P812R in Spike. Both mutations had similar patterns of change in frequency and constituted the majority of sequences by P3. Like PHE, we also detected minor variants (9), including G1251V and S1252C in Spike.

### Passage of SARS-CoV-2 on primary human airway cultures

We next sought to investigate the effect of host cell type on subsequent viral evolution, as our previous analysis assessed the contribution of viral background to viral evolution in Vero-derived cells. To this end, in parallel, we passaged SARS-CoV-2 samples on well-differentiated primary human airway epithelial cell cultures until P4, in a similar protocol as was carried out in Vero cells (**Fig 3A**). Primary cultures included WD-PNECs derived from two paediatric donors. For both PHE and BT20.1 robust infection and passage on WD-PNECs was established. For PHE, WD-PNECs were initially infected at MOI of 0.1 and virus harvested at 2-3 dpi, using the original P1 virus material. This was repeated for BT20.1, except unlike PHE, BT20.1 was directly isolated on primary cultures from the obtained clinical material. SARS-CoV-2 grew well in the primary cultures, reaching titres of ~10^6^ pfu/mL in 2-3 days in the apical compartment. Samples at each passage were subjected to sequencing as outlined above and analysed in a similar manner to those from the Vero cell passage series. For BT20.1 only P2, P3 and P4 were sequenced to compare with the data available for the equivalent Vero passage series.

**Figure 3.**
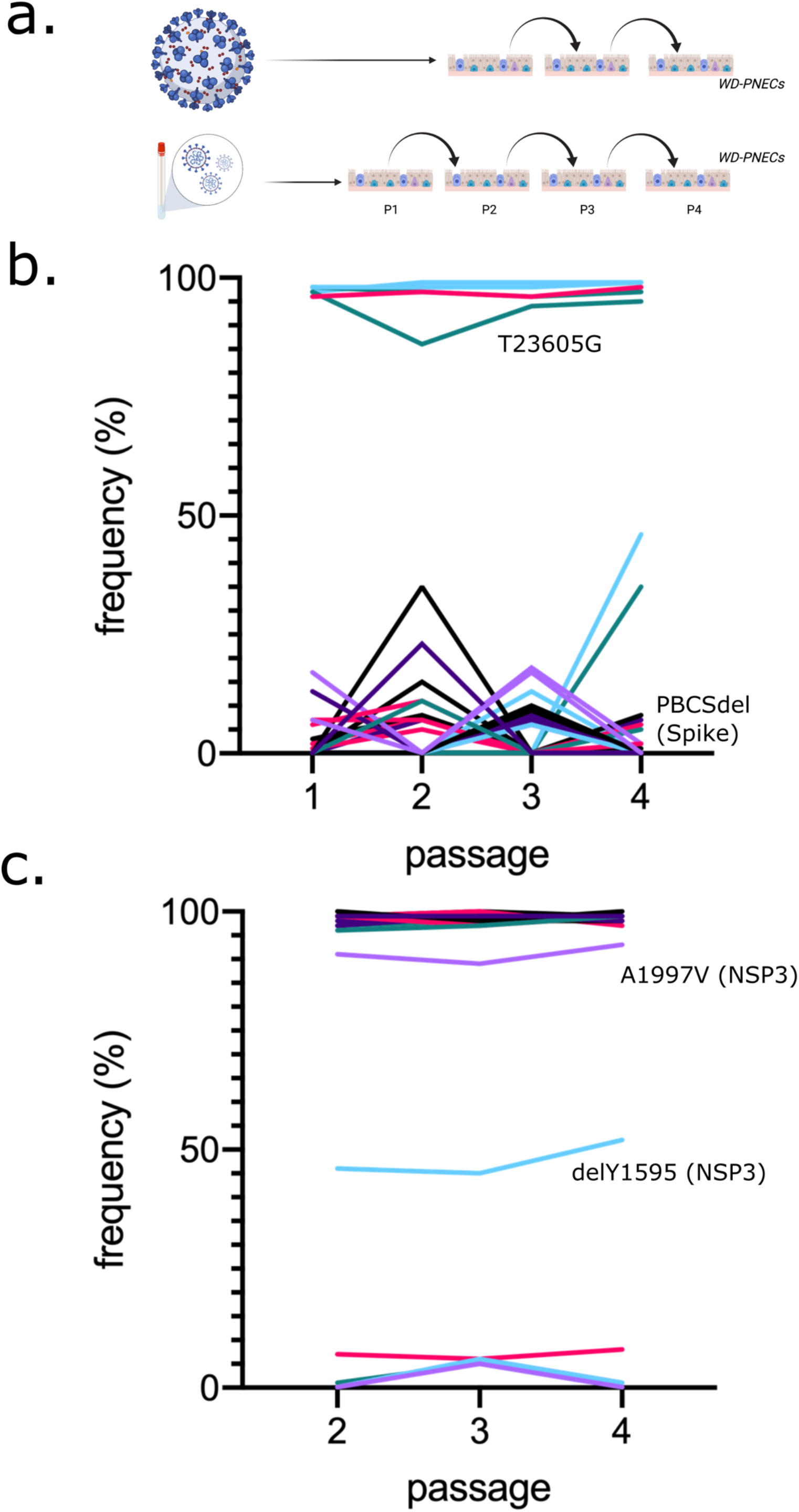
Analysis of PHE and BT20.1 whole genome sequences during WD-PNECs passage. Schematic of SARS-CoV-2 isolation/passage series on WD-PNECs for PHE and BT20.1 (a). Frequency of mutations detected for PHE (b) and BT20.1 (c) passage series on WD-PNECs, respectively, relative to the reference sequence (Wuhan-Hu-1). Only sequences from P1-P4 (PHE) and P2-P4 (BT20.1) were analysed. PHE P1 is the original stock material obtained and hence is the same sequence as PHE P1 in figure 2. Core changes were found consistently at high frequency and minor variants found at consistently low frequency (e.g. <50%). Only variants that significantly changed in frequency are marked on the graph. Colours do not reflect relationships between variants. Figures were generated with the aid of Biorender (https://biorender.com/).

In contrast to what was observed in VeroE6 cells, we did not detect any major genetic changes in PHE following passage in WD-PNECs (**Fig 3B**). However, we did identify the PBCS deletion at low levels in minor variant analysis, but never reaching majority. Together with PBCS we found 34 changes as minor variants. From passage to passage, these mutations appeared, and disappeared, stochastically. Similarly, in BT20.1, unlike the Vero cell passage, we did not find corresponding mutations in Nsp12 or Spike (**Fig 3C**). However, we identified a single amino acid deletion, Y1595, in NSP3. Intriguingly, this variant was maintained throughout the passage series at a moderate frequency of ~45%. At each passage, where possible, SARS-CoV-2 was titrated by plaque assay on Vero cells (**sFig 2**). However, we were unable to obtain titres for PHE and BT20.1 passage P3 and P1, respectively. We noticed slightly reduced titres of BT20.1 in primary cells at P4 compared to earlier passages, which was not observed for PHE (**sFig2**). WD-PNEC-grown viruses had less obvious plaques (**sFig 3 A and B)** and no evidence of cell-to-cell fusion was identified, even for BT20.1 (**sFig 3 C**). Similar to passage in Vero cells we identified two mutations in Spike (G1251V and S1252C), which appeared at low frequencies (<10%) and never increased (**table 1**).

### Phenotypic differences between SARS-CoV-2 ‘PHE’ and ‘BT20.1’ P4

Our data showing host cell dependency in viral evolution suggested differential fitness for specific viral genotypes (e.g., Vero cell-derived mutations that were not observed in WD-PNECs were less fit in primary cells). To test this hypothesis, we focused subsequent analysis on PHE and BT20.1 Vero P4 stocks with clear genetic differences between them, including the PHE PBCS deletion in Spike, and the P812R (Spike) and NSP12 mutations in BT20.1. To this end, we wished to directly compare the growth and multi-cycle replication kinetics of both strains in cell culture models of infection. To achieve this, we carried out a comparison of growth kinetics in several cell culture models, including Vero cells, VeroE6 cells modified to express human ACE2 and TMPRSS2 (VAT) (Rihn et al., 2021), and WD-PNECs (adult nasal) (**Fig 4A-C**). Of note, unmodified Vero and VeroE6 cells do not express human ACE2 and have very low levels of TMPRSS2 (**Matsuyama et al., 2020**). In Vero cells, SARS-CoV-2 grew to peak extracellular infectivity titres by ~48 hpi with titres of ~10^6^ pfu/mL. We noticed a growth attenuation of BT20.1 in Vero cells compared to PHE (**Fig 4A**). Comparing virus growth in VAT cells (**Fig 4B**), both viruses grew better but the relative attenuation of BT20.1 was not observed in VAT cells. In contrast to previous Vero cell experiments, we observed a prominent growth defect of PHE compared to BT20.1 at early time points during infection (24/48 h) in WD-PNECs. However, both viruses reached similar titres by 72 hpi (**Fig 4C**). Together these data clearly demonstrate phenotypic differences between our Vero cell-passaged viruses, demonstrating a critical role for the PBCS for efficient replication in primary cells.

**Figure 4.**
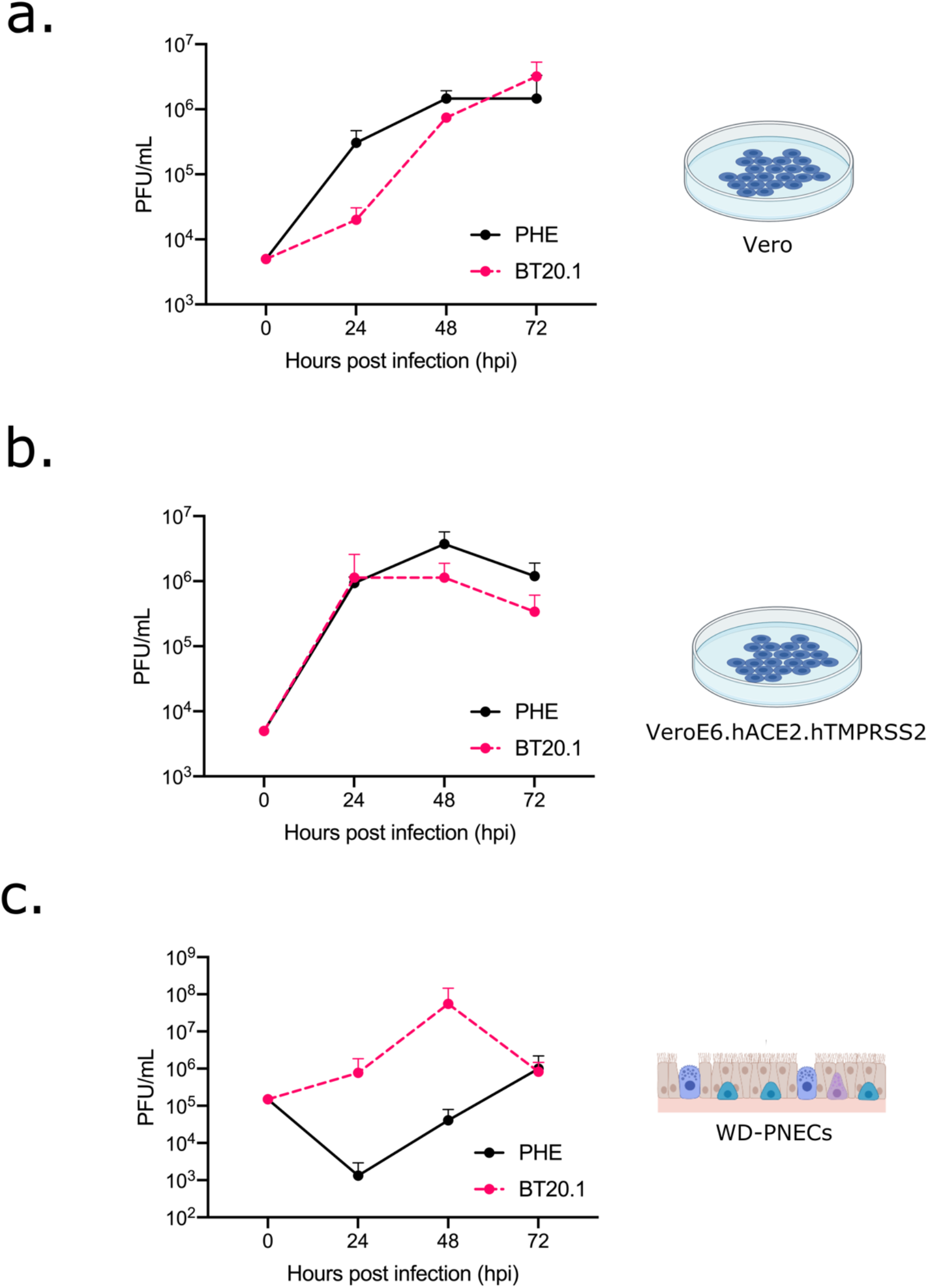
Comparison of PHE P4 (Vero) and BT20.1 P4 (Vero) growth on different cell substrates. Multicycle growth curves (MOI 0.01 for Vero or 0.1 for WD-PNECs) for PHE P4 (VeroE6) and BT20.1 P4 (Vero) on Vero (a), VeroE6 cells expressing human ACE2 and human TMPRSS2 (b), and adult WD-PNECs from 3 donors (c). Titres for Vero-derived cells are shown as means +/- SEM for triplicate wells and are representative of two independent experiments. Titres for WD-PNECs are shown as means +/- SEM for single wells from 3 donors. Data using BT20.1 are presented here as averages from 3 donors but have also been incorporated into a sister paper using separated, individual donor data (Broadbent et al., 2021 in submission). Figures were generated with the aid of Biorender (https://biorender.com/).

## Discussion

Investigation of the patterns of SARS-CoV-2 genetic diversity worldwide during outbreaks has already facilitated a genetic-based nomenclature of lineages and has also highlighted the emergence of functionally-relevant mutations, such as D614G in Spike, and those contained in extant VOCs (**Harvey et al., 2021**). Complementary to this, in vitro systems are an incredibly useful and tractable means to understand the forces influencing this viral evolution, in particular those that mimic in vivo-relevant conditions, such as WD-PNECs (**Hou et al., 2020**).

Our data and that of others demonstrate significant standing genetic diversity in viral populations in vitro that can be acted upon by rapid evolutionary processes. Although also observed in our work, from early in the study of SARS-CoV-2 evolution in Vero-like cells, it was revealed that the virus readily diversifies during culture with the most evident being mutations mapping to the PBCS of Spike (**Davidson et al., 2020; Klimstra et al., 2020; Lamers et al., 2021; Ogando et al., 2020; Pohl et al., 2021**). Consistent with our work, other studies have identified enhanced genetic stability, in particular of the PBCS only, during passage of one strain in Calu3 or primary airway organoids (**Lamers et al., 2021**). One striking finding of our work, which builds on previous studies, is that on several occasions for both isolates we observed a rapid increase in frequency of specific mutations in the PBCS and independently of it (PBCS deletion & P812R in Spike, and T293I in NSP12) to near fixation over the course of a couple of passages in Vero cells. These patterns suggest a selective phenotypic advantage in that particular cell culture system. Similar changes (including P812R and the NTD deletion) were identified in other studies (**Dieterle et al., 2020; Ramirez et al., 2021**). The fact that identical mutations arise independently (e.g., loss of the PBCS and P812R) is highly suggestive of convergent evolution, perhaps toward a similar phenotype. Our work on the loss of the PBCS in Vero cells and its association with attenuation in WD-PNECs is consistent with previous reports. However, it is noteworthy that we did not observe deletion or mutations in and around the PBCS during passage of BT20.1 in Vero cells.

In addition to the loss of the PBCS, we observed P812R in Spike and T293I in NSP12, although we were not able to associate them with changes in virus growth in WD-PNECs due to a lack of an additional comparable ‘wild-type’ isolates. However, the fact that parallel passage in WD-PNECs did not result in their increased frequency suggests that they confer a hitherto unrecognised disadvantage in the primary epithelial cell system. P812R is a non-conservative change and rapidly rose to near-fixation alongside NSP12 in BT20.1 in Vero cells. P812 sits near the S2’ cleavage site and is a highly conserved position in SARS-CoV-2. However, non-P residues (e.g. serine) were occasionally found in nature but are rare (https://nextstrain.org/), which suggests a functional defect in vivo. Interestingly, P812R was observed before, in at least two other studies, associated with a change in Spike activity using full-length SARS-CoV-2 and one using chimeric vesicular stomatitis virus encoding SARS-CoV-2 Spike (**Dieterle et al., 2020; Ramirez et al., 2021**). Like previous work, we noted an association of P812R with enhanced cell-to-cell fusion when BT20.1 grown on Vero cells is compared to that grown in WD-PNECs (**sFig 3**). It was suggested that P812R generated a novel PBCS at the S2’ site (**Ramirez et al., 2021**). Cleavage by furin-like proteases could thus compensate for lack of TMPRSS2-mediated proteolysis and activation in Vero cells. Although it is possible that P812R confers a similar phenotypic change as the PBCS deletion, it is not likely to be identical, given the clear differences in growth between PHE and BT20.1 in Vero cells and WD-PNECs. Along with P812R in S, BT20.1 carried a mutation in NSP12 (T4685I/T293I), which is the viral RNA-dependent RNA polymerase. The mutation sits on the surface in close proximity to a zinc-binding site of the interface domain that mediates intra-NSP12 interactions and interactions between NSP12 and other polymerase co-factors, such as NSP8 (**Hillen et al., 2020**). Given the linkage between P812R and T4685I, further molecular virological work using isogenic viruses generated through reverse genetics is required to ascertain the impact of this mutation in relevant cell models. The mutations T4685I arose with P812R possibly suggesting genetic linkage, although this remains to be determined.

It is of interest that BT20.1 carries a deletion in ORF7A that results in a frameshift and C-terminal truncation of the protein, likely ablating the transmembrane domain and tail. ORF7A is a type 1 transmembrane protein and it has numerous putative functions involved in host-pathogen interactions and immune evasion (**Nemudryi et al., 2021**). ORF7A truncations in SARS-CoV-2 isolates have been discovered before, possibly associated with reduced capacity to subvert the innate immune response (**Nemudryi et al., 2021**). However, the previous work was carried out using non-clinically relevant cell models, such as Vero or HEK-derived lines. Our work suggests that full-length ORF7A is not required for replication in Vero or WD-PNECs and likely serves an accessory function that may affect replication and/or transmission in particular circumstances.

Not only did we observe changes reaching near-fixation in our dataset we also identified several lower frequency mutations in our viral populations. Consistent with this variation within a stock, we also noticed plaque size variation in passage stocks suggestive of functional differences between viral sub-clones (**Fig 1 E and F**). We detected an in-frame deletion of a single codon in the C-terminus of NSP3, located in the Y1 domain, which is located on the cytoplasmic face of the virus-remodelled ER membrane, where it may regulate replication complex stability by interacting with NSP4 (**Lei et al., 2018**). NSP3 itself is a multifunctional protein involved in numerous viral processes. The fact that the deletion did not rise to fixation suggests that it is at a competitive disadvantage compared to wild-type. The mutation in NSP3 is also interesting because it is maintained at a moderate frequency. Of considerable interest is the overlap between variations observed in Vero cells and that of VOCs, especially in the NTD of Spike. We observed three mutations in the NTD in PHE P3 and P4: E180K, D215G, and a deletion resulting in the loss of 9 amino acids. Variants identified in this study mapping to the ectodomain of Spike are marked on a structural model (**sFig 4**). For D215G and the deletion, these mutations are similar to those in VOCs, such as alpha and beta variants. Regarding mutations in NTD loops, several VOCs have convergently modified the amino acid identity of the loop. While in vivo this may be the result of antibody selection, in our system there are no antibodies, which suggests a role for NTD mutations independent of antibody selection. The rise in frequency is suggestive of a fitness advantage of these mutations. Further work is required to determine the function of the NTD of Spike and the impact of these mutations on the virus life cycle.

While general trends were similar between our two isolates in Vero cells (i.e., mutations rising to high frequency), specific mutations observed were not. It is likely that evolution of key mutations reflect inherent biological differences in viruses and not subtle changes in passaging conditions. In PHE, loss of the PBCS occurred, which was not observed in BT20.1, and vice versa regarding P812R and NSP12. This is consistent with an effect dependent on viral input or strain or genetic background through epistatic interactions between mutations, such as D614G in Spike. However, in numerous reports, isolation and passage of SARS-CoV-2 on Vero cells selected for a loss of the PBCS, which was not observed in our BT20.1 passage series. Alternatively, the mutation P812R could functionally achieve the same phenotype as the deletion of the PBCS, although our primary cell infection model where PHE was attenuated compared to BT20.1, would suggest that this is not the case.

By comparing evolution of the same isolates in two distinct cell culture systems, we observed a dependence on host cell substrate on downstream virus evolution. Namely, passage in WD-PNECs resulted in enhanced stability of SARS-CoV-2 genetic diversity at the consensus level. While the PBCS, P812R and NSP12 changes were identified in PHE and BT20.1 when grown in Vero cells, these changes did not rise to high frequencies in WD-PNECs. Differential accumulation of mutations may reflect distinct host cellular environments encountered upon passage in Vero or WD-PNECs. This reflects major differences in these cell substrates, including i) species and tissue differences; ii) reduced levels of TMPRSS2 in Vero cells; iii) reduced innate immune response in Vero cells as they are deficient in type 1 interferon production (**Emeny & Morgan, 1979**). However, what affects the rise in P812R/NSP12 mutation remains unknown. Future work will assess the effect of these changes in BT20.1 upon replication in WD-PNECS. Additionally, during passage of BT20.1 in WD-PNECs we identified a deletion in Nsp3, although the relevance and mechanism of this change is unknown.

In conclusion, by studying the evolution of SARS-CoV-2 during passage in distinct cellular substrates we shed light on the forces that shape viral fitness, unveiling a collaboration between both viral and host factors in driving SARS-CoV-2 genetic diversity, which helps define the molecular correlates of fitness in the natural target cells. Finally, on a practical note, our results support close characterisation of virus stocks for experimentation in vitro and in vivo and suggest ways to mitigate unwanted cell culture artefacts, critical for understanding host-pathogen interactions and identification of antiviral interventions.

---

**sTable 1.**
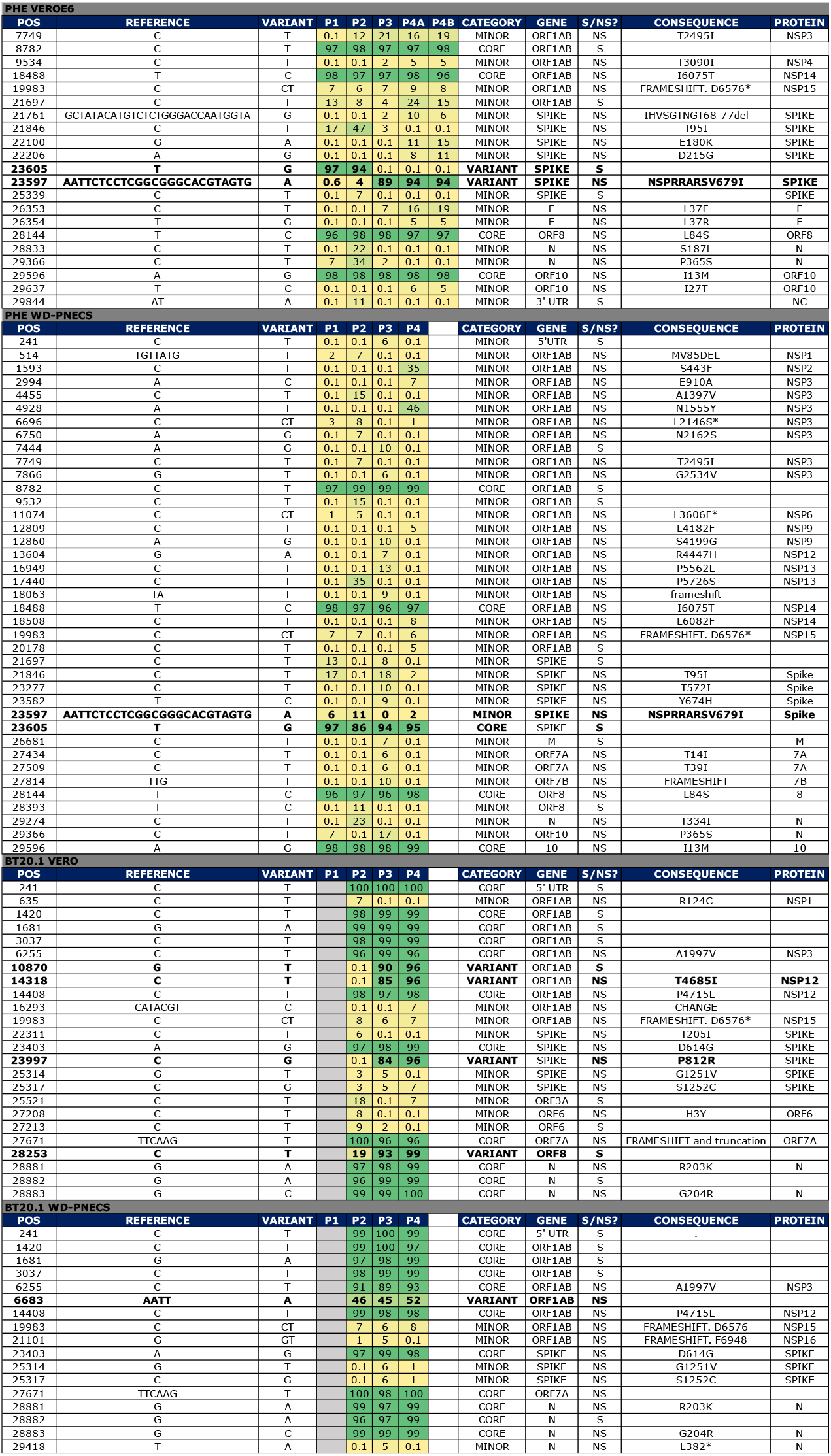
Frequency of variants in reference to Wuhan-Hu-1 identified in this study. Variants only shown where there was at least one instance of frequency >5%. Where undetectable an arbitrary value of 0.1 was assigned. Frequency data is highlighted by colour (green for higher, yellow for lower). Mutations have been assigned status of core, variant or minor. Additionally, for each variant, data for nucleotide location, reference and variant nucleotides, gene & protein location, and consequence (e.g., synonymous [S] or non-synonymous [NS]) are shown.

**sFig 1.**
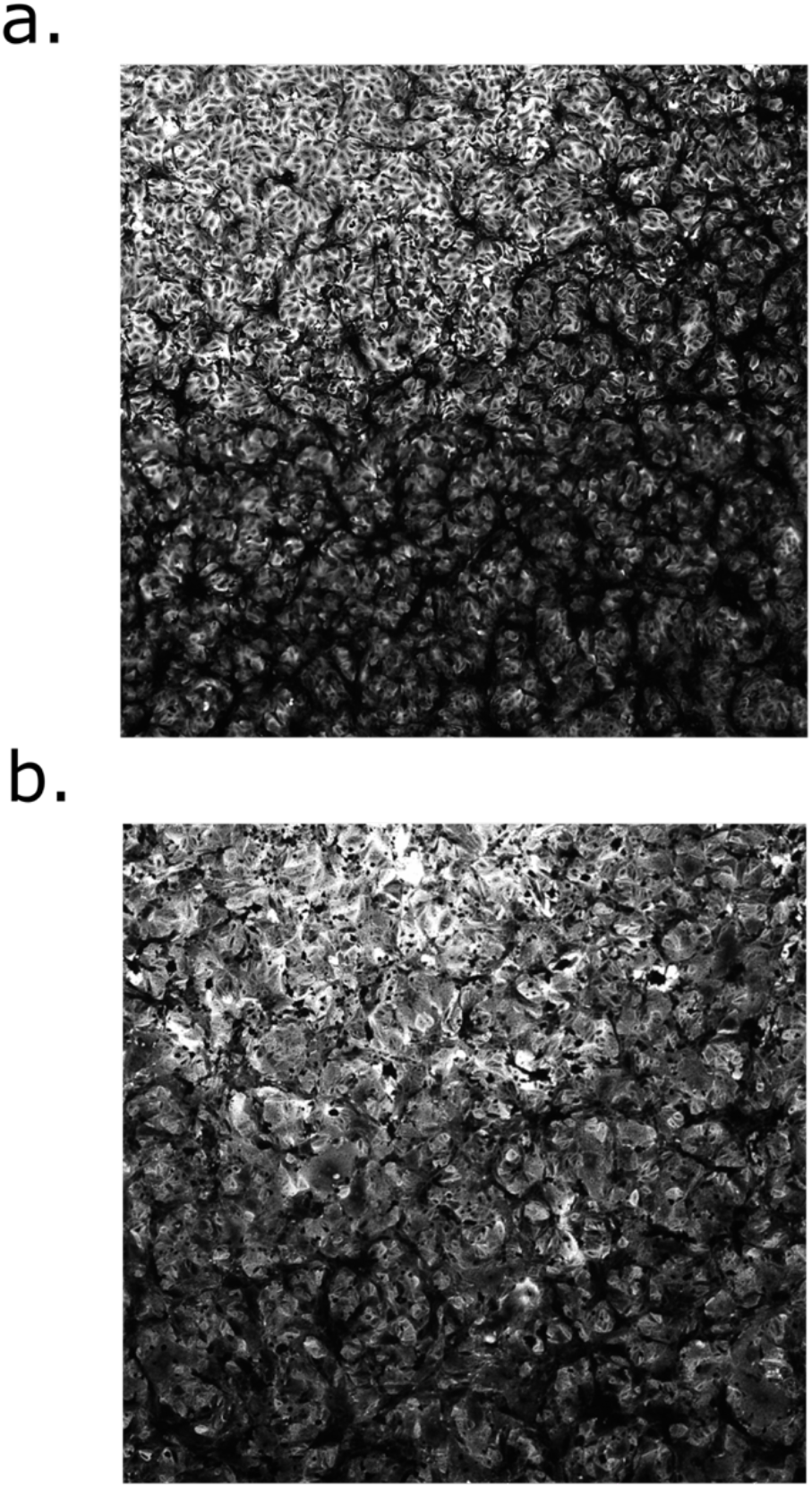
Fusogenicity of PHE and BT20.1 P4 on Vero cells. Higher magnification images of plaque visualisation of PHE (a) and BT20.1 (b) P4 on Vero cells from the same images shown in Figure 1.

**sFig 2.**
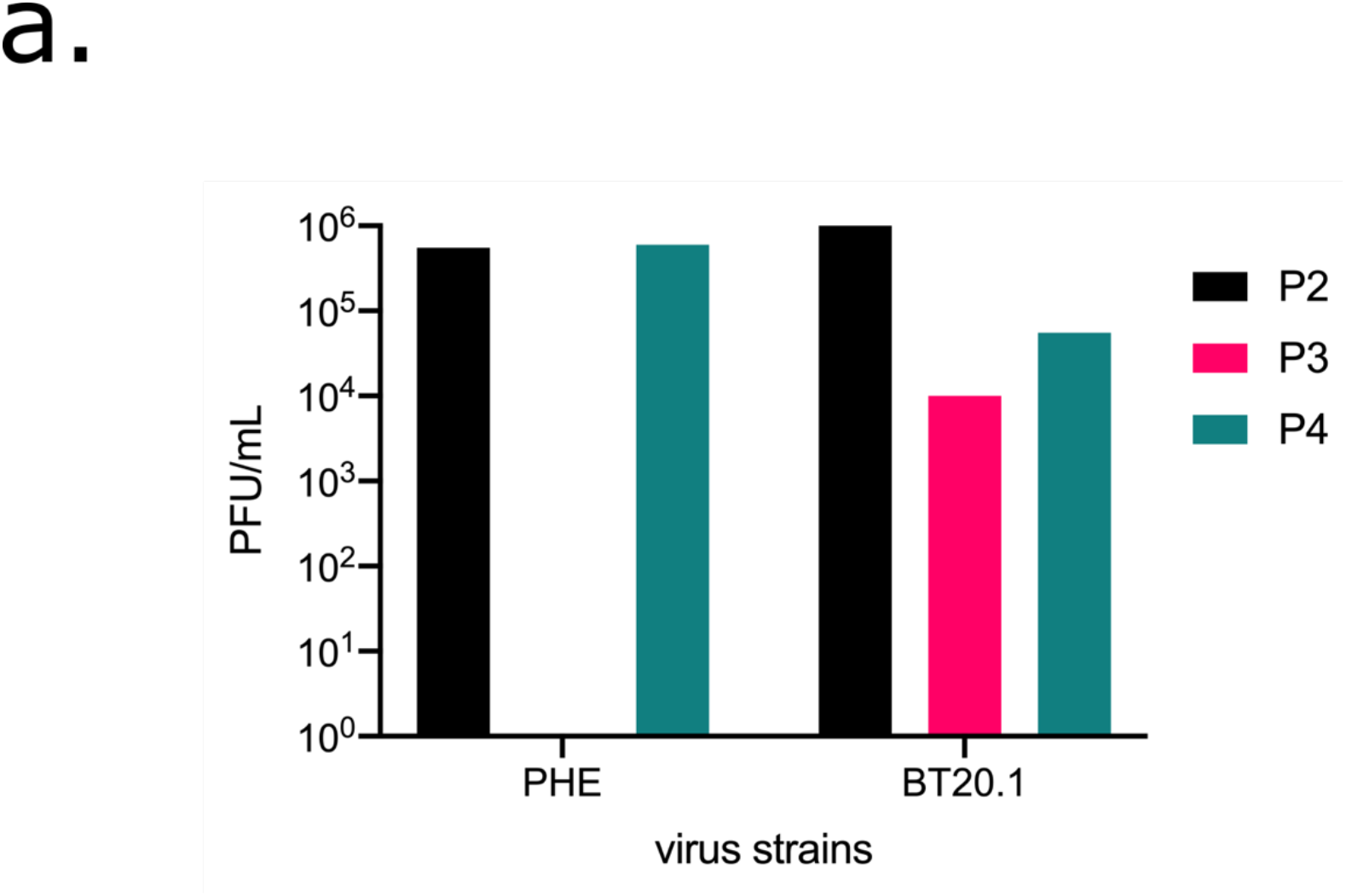
Growth kinetics of PHE and BT20.1 during passage in WD-PNECs. Infectivity titres for material generated from isolation/passage of PHE and BT20.1 on WD-PNECs.

**sFig 3.**
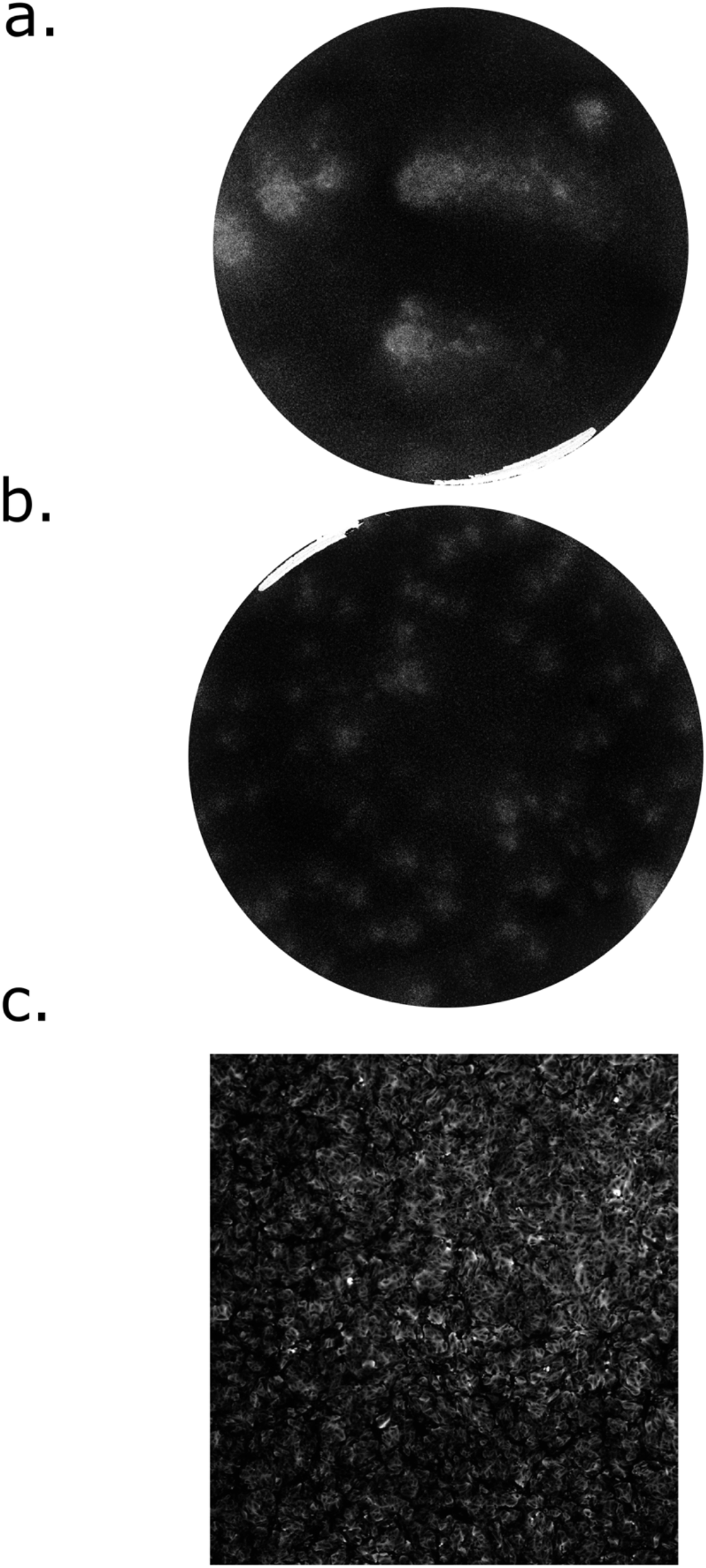
Plaque morphology of SARS-CoV-2 grown in WD-PNECs. Plaque visualisation of PHE (a) and BT20.1 (b) P4 on Vero cells. Higher magnification images of plaque visualisation of BT20.1 (c) P4 on Vero cells from the same images shown (b).

**sFig 4.**
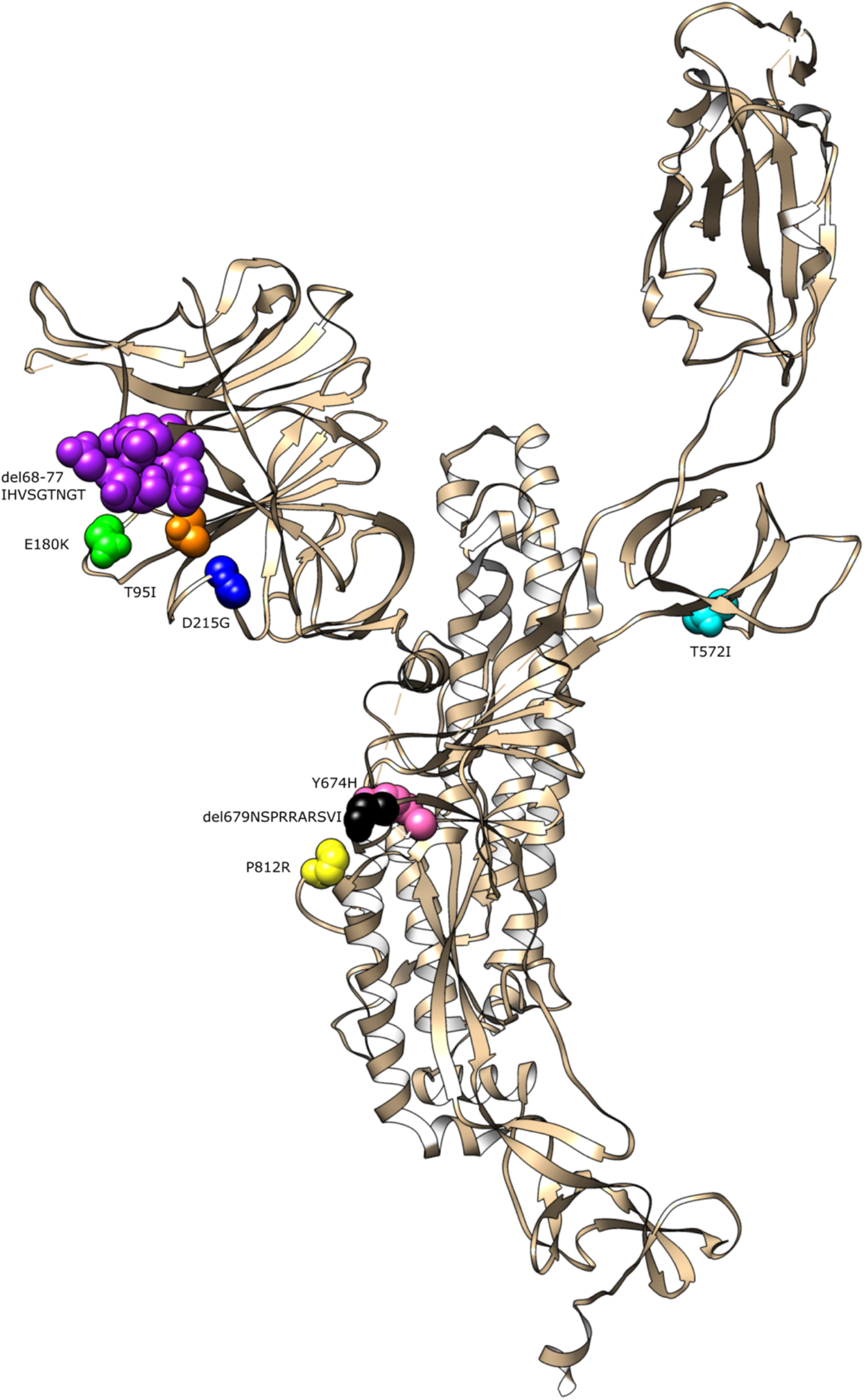
Location of Spike mutant variants observed in this study on model structure of a single Spike monomer in the pre-fusion state (PDB 7C2L from (Chi et al., 2020)). Variants identified in the Spike cytoplasmic tail (G1251V and S1252C) are not shown.

## Materials and methods

### Continuous cell line culture

In this study, 3 continuous cell lines were used: Vero wildtype (number), Vero E6, and Vero E6 expressing human ACE2 and TMPRSS2 (VAT) (Rihn et al., 2021). All cells were grown in DMEM (5% FCS v/v) with antibiotics. VAT cells were maintained in the presence of additional antibiotics to select of cells carrying transgenes. Cell lines were routinely tested for mycoplasma contamination and no evidence of contamination was detected.

### WD-PNECs

Nasal epithelial cells from preschool age children with recurrent wheeze (for initial passaging) and from healthy adults (for final comparison of PHE and BT20.1 P4 viruses) were obtained by brushing of the nasal turbinates with an interdental brush (DentoCare). Cells were cultured in monolayer until passage 3 then seeded onto collagen-coated Transwells (6 mm, 0.4 μm pore size; Corning). Once confluent the apical medium was removed to create an air-liquid interface which, together with specialised media (Pneumacult ALI, Stemcell Technologies), triggered differentiation (Broadbent et al. 2016. Broadbent et al. 2020). Complete differentiation (after a minimum of 21 d) was confirmed by an intact culture, extensive cilia coverage and mucus production.

### Viruses

Two SARS-CoV-2 isolates were used throughout this study, including ‘PHE’ and ‘BT20.1’. PHE was provided as an early passage isolate on VeroE6 cells while BT20.1 was provided directly as a nasopharyngeal swab in virus transport media clinical material from a positive case from Belfast in June 2020. Stocks were prepared in Vero or VeroE6 cells in DMEM containing 2.5% FCS (v/v) infected at a low MOI (~0.001). Infections were harvested when maximal cytopathic effect was noted, usually between 3-4 days post infection. Infected culture supernatant was harvested, clarified by centrifugation and stored at −80°C. WD-PNECs were apically infected with SARS-CoV-2 for 1 h, after which the inoculum was removed and the apical surface gently rinsed with DMEM. Virus was harvested from WD-PNECs by incubation of the apical surface with DMEM for 5 min at room temperature in the absence of serum. Harvested virus was immediately stored at −80. All SARS-CoV-2 work was carried out under BSL3 conditions in a dedicated facility in QUB.

### Plaque assays

Our plaque assay protocol is based on the methodology available here: https://www.protocols.io/view/viral-titration-of-sars-cov-2-by-plaque-assay-semi-be4zjgx6. Near confluent monolayers of Vero cells in 24 or 6 well plates were infected. On the day of titration growth media was replaced with DMEM (0% FCS) (250 μL). Virus dilutions were prepared in plate and incubated for 30 min after which the 2x overlay medium (containing 2% agarose) was added. Plates were incubated for 3 days at 37°C. At 3 dpi PFA (8%) was added to the cultures and cells fixed/inactivated for at least 20 min. Following fixation, the PFA was removed and monolayers stained for 10 min using crystal violet (1% w/v in ethanol 20%). Following staining, residual crystal violet solution was removed, plates were rinsed in water and submerged in Chemgene prior to drying and removing from the hood for visualisation and quantification. To calculate PFU/mL, plaques at a dilution were quantified, the precipice of this number used, and multiplied by the dilution factor (4). For visualisation of plaque assays, whole plates were scanned using a Celigo imaging cytometer (Nexcelom Bioscience).

### Virus whole genome sequencing

Virus whole genome sequencing used methods developed by the ARTIC network (https://artic.network;(**Tyson et al., 2020**)) and the COG-UK Consortium. Culture supernatants were inactivated by addition of Triton X-100 to 1.5% (v/v). Viral RNA (total nucleic acid) was extracted from inactivated samples (200 μL) using the MagNA Pure Compact instrument and MagNA Pure Compact Nucleic Acid Isolation Kit I (Roche Molecular Systems Inc, Burgess Hill, UK). Purified nucleic acid was eluted into 100 μL and used immediately or stored at −80°C. For first-strand cDNA synthesis, nucleic acid (5 μL) was used as template for reverse transcription using LunaScript® RT SuperMix Kit (New England Biolabs, Hitchin, UK) in 20 μL reaction volume. Primers were annealed (65°C, 5min, snap-cool on ice) prior to addition of reverse transcriptase. Reactions were incubated at 42°C (50 min) then stopped at 70°C (10 min). The resulting cDNA was used immediately for PCR or stored at −20°C. In brief, these were run as two separate multiplex PCR “pools” (A & B) using the ARTIC version 3 primer set (ARTIC nCoV-2019 V3 Panel, IDT DNA Inc, Leuven, Belgium; https://github.com/artic-network/primer-schemes/tree/master/nCoV-2019) and Q5 DNA polymerase mastermix (New England Biolabs). Following PCR, the amplicons from pools A & B were combined, and the resulting pooled amplicons (98 x 450 bp overlapping tiled amplicons, spanning the SARS-CoV-2 genome) were purified using Kapa HyperPure beads (Roche Molecular Systems Inc) and quantified using a Qubit fluorometer and dsDNA HS Assay Kit (Thermo Fisher Inc, Manchester, UK). Amplicon sequencing libraries were prepared using the Nextera DNA Flex library kit according to the manufacturer’s instructions (Illumina Ltd., Cambridge, UK). Libraries were sequenced on a MiSeq (Illumina) using a MiSeq Reagent Kit v2 and 2 x 151 bp paired-end sequencing protocol (Illumina).

### Sequence analysis

The FASTQ files were uploaded into the Galaxy web platform, and we used the public server at usegalaxy.eu to analyse the data (**Afgan et al., 2018).** The workflow used was specially optimized for Illumina-sequenced based ARTIC pair end data with the intention to detect allelic variants (AVs) in SARS-CoV-2 genomes (**Maier et al., 2021**). This analysis converted FASTQ data to annotated AVs through a series of steps that include QC, trimming ARTIC primer sequences off reads with the iVar package, mapping using bwa-mem, deduplication, AV calling using lofreq, and filtering AVs that both occurred at an allele frequency (AF) ≥5%, and were supported by ≥10 reads. As we could not determine the background frequency of mutations we focused on those variants with a minor allele frequency ≥5%, and were supported by ≥10 reads in at least one passage of the series. Furthermore, we focused our greater analysis on those found in more than one passage and those that substantially rise in frequency. Raw data and consensus sequences will be uploaded during review and before publishing.

## Acknowledgements

The authors would like to thank the following individuals and organisations for provision of critical reagents, and technical assistance: Public Health England and Professor Maria Zambon for the original SARS-CoV-2 England 02/20 isolate; Belfast Health and Social Care Trust for the original SARS-CoV-2 clinical material from which BT20.1 was derived; Professor Alain Kohl (MRC-University of Glasgow Centre for Virus Research) for the VeroE6 cells; and Dr Suzannah Rihn (MRC-University of Glasgow Centre for Virus Research) for the VeroE6 cells expressing human ACE2 and human TMPRSS2; QUB staff Mervyn McCaigue, Cathy Fenning, David Norwood, Nuala McCann, Paul Crowe and Zoe Hunter for their assistance in establishing and maintaining the QUB SARS-CoV-2 BSL3 facility; and finally, Professor Jose Bengoechea for unwavering support for QUB virology at this critical time. We also wish to thank the QUB Genomics Core Technology Unit for their help with sequencing. Funding from UKRI/NIHR (MC**_** PC**_**19057) to UP, KM; PHA HSCNI R&D Division (COM/5613/20) to UP, KM, LB, CGGB, and GLC; and generous donations from the public to the Queen’s University of Belfast Foundation was used for the study.

